# Climate change is predicted to simplify seed dispersal networks in the Cerrado

**DOI:** 10.64898/2026.04.30.721967

**Authors:** Eduardo D. B. Rigacci, Mariana Campagnoli, Jeferson Vizentin-Bugoni, Alexander V. Christianini, Guadalupe Peralta

## Abstract

1. Animal-mediated seed dispersal is key for the maintenance and functioning of tropical ecosystems. Specifically, in the Cerrado, the largest Neotropical savanna and a global biodiversity hotspot, nearly 60% of plant species rely on animals for dispersal.
2. Climate change threatens these interactions by affecting species distributions, reshaping communities, and potentially decoupling plants from their dispersers. Anticipating how such disruptions may alter seed dispersal networks is particularly relevant for understanding the resilience of future tropical ecosystems.
3. Here, we combined empirical data on 139 pairwise plant-frugivore interactions with species distribution forecasts to build probabilistic interaction matrices under present and future climate scenarios, which were then used to construct 6,221 local seed dispersal networks. Using ecological niche modelling, we tested how climate change influences species’ range size and centroid displacement. Then, we evaluated whether such changes translate into losses of pairwise plant-frugivore co-occurrence. Finally, we investigated how these changes in occurrence overlap may affect key structural properties of future local seed dispersal networks.
4. We forecast that by the 2070s, under a business-as-usual climate scenario, species are likely to contract their ranges by 56 ± 33% and shift their distribution centroids by 88 ± 57 km within the Cerrado, leading to a 27 ± 29% loss in plant-frugivore co-occurrence mainly driven by reductions in plant species distributions. At the community level, these losses will lead to smaller and more nested networks and specialized, indicating a structural simplification of seed dispersal systems in the Cerrado.
5. Synthesis: By combining empirical data on animal-mediated seed dispersal with forecasts of species distributions, we found that climate change may simplify frugivore-plant interaction networks in the Cerrado by decreasing species ranges and co-occurrence of partners. Our study demonstrates that future climate may pose a threat not only to species distributions but also to ecological interactions, such as seed dispersal, that are key to enabling climate-tracking by plants. Thus, preventing the simplification of interaction networks will be essential to conserve biodiversity in species-rich regions.

## 1. INTRODUCTION

In tropical ecosystems, animal-mediated seed dispersal is a cornerstone ecological process that shapes community dynamics, where many vertebrates include fruits in their diet and the majority of woody plants depend on animal dispersers (Jordano, 2000). These mutualistic interactions allow plants to colonize new sites, maintain metapopulation connectivity, and reduce intraspecific competition (Comita et al., 2014; Jordano, 2000). In return, frugivores obtain water, nutrients, and energy from fruit pulp and arils (Howe & Smallwood, 1982). The structure of seed-dispersal networks can be influenced by multiple processes (Campagnoli, Christianini, et al., 2025; Vizentin-Bugoni et al., 2022). For instance, morphological trait matching, such as the match between fruit size and bird gape width, can constrain interaction occurrence and influence the number of seeds dispersed per visit (Campagnoli, Christianini, et al., 2025; Dehling et al., 2014). Similarly, geographical co-occurrence and phenological overlap between plants and frugivores can substantially influence both interaction occurrence and frequency (Campagnoli, Christianini, et al., 2025; Vizentin-Bugoni et al., 2022). However, by reshaping species distributions, climate change may create spatiotemporal mismatches between currently interacting species (Sales, Culot, et al., 2020; Schleuning et al., 2020).

Changes in temperature and precipitation under future climates may lead to contractions or expansions of species’ geographic ranges (Nascimento et al., 2025; Velazco et al., 2019), depending on species attributes such as physiological tolerances, life-history strategies, and dispersal capacities (Parmesan, 2006). Specifically, species can span a gradient of environmental change tolerance, from highly tolerant (e.g., habitat generalists, wide-ranging and synanthropic species), which are expected to be less affected by climate change and may even expand their ranges, to highly sensitive (e.g., habitat specialists and range-restricted species), which may become locally extinct due to the loss of suitable habitats (Filgueiras et al., 2021). Therefore, species-specific responses to novel climatic conditions may result in asynchronous range shifts among interacting animals and plants, leading to spatial decoupling of seed-dispersal interaction partners.

Climate-driven loss of species co-occurrence may substantially affect the structure of seed dispersal networks. For instance, climate change is projected to reduce the number of frugivore species locally (Mota et al., 2022), with trophic and habitat specialist taxa potentially being more susceptible to such changes (Gonçalves et al., 2021). Thus, networks are likely not only to shrink but also to become dominated by generalist species, as observed under other environmental change drivers (Li et al., 2022). The persistence of a highly connected core of generalist taxa may erode the hierarchical organization of interactions, yielding networks that are structurally simplified, which may ultimately affect ecosystem function and resilience (Li et al., 2022).

Despite the importance of understanding the effects of climate change on seed dispersal networks, the subject remains poorly explored in many climate-vulnerable biodiversity hotspots, such as tropical grassy biomes (Goel et al., 2020), which cover up to approximately 40% of Earth’s terrestrial surface (Parr et al., 2014). Particularly, in the Cerrado, a megadiverse tropical savanna where about 64% of woody plant species depend on animals for dispersal (Gottsberger & Silberbauer-Gottsberger, 1983), ongoing climate change has already extended and intensified the dry season and increased the risk of anthropogenic wildfires (Hofmann et al., 2023). While some studies indicate climate change may reduce the range size of frugivore (Hidasi-Neto et al., 2019, 2022) and plant species’ (Simon et al., 2013) in the Cerrado, the potential effects of these changes on species co-occurrence and their interactions remain largely unknown.

Here, we asked whether climate change will affect plant-frugivore seed dispersal interactions in the Brazilian Cerrado. Specifically, we tested how climate change influences species’ range size and range displacement, and whether such changes translate into losses of pairwise plant-frugivore co-occurrence. Then, we assessed how these co-occurrences changes affect key structural properties of seed dispersal networks, including species and interaction richness, weighted nestedness, and specialization. We expect to find a forecast indicating overall reduction of plants and animals ranges, decreasing partners’ occurrences and, consequently, smaller and more simplified interaction networks.

## 2. METHODS

In short, we combined empirical data on 139 pairwise plant-frugivore interactions known to occur and interact in the Cerrado (see Campagnoli et al., 2025). We estimated current plant and frugivore species distribution to assess whether different climate change scenarios affected species range size and displacement. We then combined species distribution information with species temporal overlap and trait matching to predict changes in interaction patterns at local ecological networks due to climate change (Figure 1).

**Figure 1.**
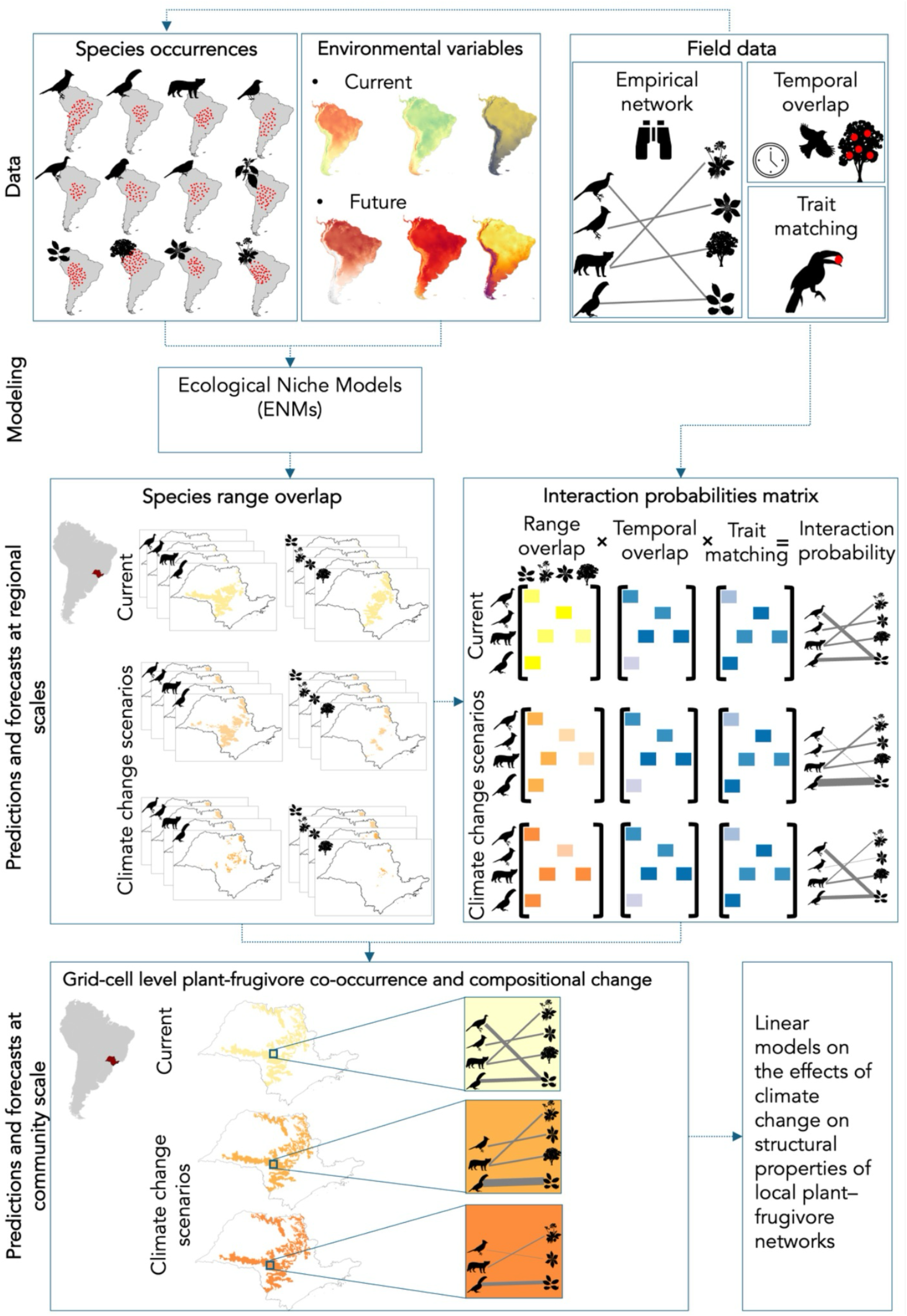
Workflow of data integration, modelling, and statistical analyses used to assess the effects of climate change on plant–frugivore interaction networks in the Cerrado. Species occurrence records and environmental predictors were compiled from multiple sources and used to fit ecological niche models (ENMs), which were then projected under current and future climate scenarios. Empirically derived plant–frugivore interaction data from (Campagnoli, et al., 2025)were used to define the study species pool and to quantify phenological overlap and trait matching between interacting partners. Current and future species distributions were used to estimate range overlap in the Cerrado of São Paulo State (indicated in red), which was combined with phenological overlap and trait matching to generate interaction probability matrices for each scenario. These matrices were then used to reconstruct local seed-dispersal networks at the grid-cell level (ca. 50 km²) based on predicted species co-occurrence. For each grid cell, we calculated descriptors of network structure and used them in statistical models to test the effects of climate change on plant–frugivore networks.

### 2.1 Study area and Dataset

Our study area encompassed the Cerrado *sensu lato* in São Paulo State, Brazil, with a spatial extent of approximately 101,000 km², which included major transitional zones with the Atlantic Forest (Durigan & Ratter, 2006). This region has been almost entirely converted to intensive agriculture, particularly for soybean and sugarcane cultivation, leading to severe defaunation (Durigan & Ratter, 2006; Strassburg et al., 2017). Despite this long history of land-use conversion, the Cerrado remnants in São Paulo still represent some of the most ecologically significant patches in its southernmost distribution (Ratter et al., 2003).

The interactions and species-level data were sampled at Itirapina ecological station (22°12′ S, 47°51′ W) at São Paulo state, Brazil, one of the largest remaining fragments of Cerrado *sensu lato* in southern Brazil. The empirical dataset used here comprises 139 links of plant–frugivore interactions involving 3 mammals, 35 birds, and 30 plant species. To our knowledge this dataset represents one of the most comprehensive seed dispersal networks for a Cerrado site. We recorded plant-frugivore interactions, fruit and frugivore phenology, and morphological traits in six 100 m x 10 m transects during January 2021 to October 2022 at Itirapina, spanning a gradient of tree cover that captures the variation in vegetation density typical of Cerrado landscapes. Plant-frugivore interactions were recorded by combining three different sampling methods: focal observations, faecal samples from birds captured with mist nets, and videos recorded using camera traps (see “Sampling of plant-frugivore interactions” in the Supplementary Material). We considered an interaction event every time a frugivore visited and removed fruits from a plant, regardless of the number of fruits consumed, or when seeds were found within birds’ faecal samples. Interactions involving animals that only consumed fruit pulp, and/or dropped all seeds beneath parental plants, were not included in our analyses as we were interested in seed dispersal interactions.

Fruit and frugivore phenology and morphological traits were also retrieved from the previously mentioned dataset. Authors recorded monthly fruit phenology through the presence of mature fruits inside each transect, and monthly frugivore phenology through the detection of birds and mammals during point counts and videos from camera traps, respectively. Gape and gullet sizes were recorded based on captured bird individuals, measurements from museum specimens at the USP Zoological Museum (MZUSP) and complemented with data from published trait databases (Bello et al., 2017; Fuzessy et al., 2022; Tobias et al., 2022; Wilman et al., 2014). Fruit diameter was measured directly in the field following standardized protocols (Pérez-Harguindeguy et al., 2013), and average values were calculated per plant species.

We used as our study area the Cerrado of São Paulo State, southeastern Brazil, mainly because we had empirical data from this region Because this region represents a distinctive southern biogeographic unit within the Cerrado, shaped by its floristic composition, environmental conditions, transition with the Atlantic Forest, and long history of land-use conversion to croplands (Durigan & Ratter, 2006; Ratter et al., 2003; Strassburg et al., 2017), its unique configuration may result in ecological filters for both plants and frugivores, influencing the composition and structure of local seed-dispersal networks. Therefore, extrapolating the empirical seed-dispersal network obtained within this biogeographic unit to the entire Cerrado would be ecologically unjustified, leading us to restrict the spatial extent of our analyses to maintain the ecological realism of our forecasts.

### 2.2 Current species distribution

To estimate plant and frugivore species distribution in the Cerrado and forecast how species may redistribute under future climate scenarios, we built Ecological Niche Models (ENMs). ENMs combine species-specific occurrence records with environmental variables to obtain species-environmental relationships and then project these relationships to the geographical space (Fig. 1) (Guisan & Zimmermann, 2000; Peterson et al., 2011). For the empirical plant-frugivore interaction dataset, we collected occurrences throughout this region from comprehensive regional datasets and global biodiversity data aggregation infrastructures (see “Species occurrence sources” in the Supplementary Material text) (Fig. 1). We then cleaned, quality-checked, and thinned the species occurrence records. This resulted in 214,874 records, averaging 3,207 ± 4,016 records per species. Among frugivores, *Pitangus sulphuratus* had the greatest number of records (32,384) and *Antilophia galeata* the fewest (247); for plants, *Myrcia guianensis* had the most records (1,532) and *Smilax polyantha* the least (95). Details on the cleaning procedures are provided in the “Data cleaning” section in the Supplementary Material text and Supplementary Table S1.

As environmental variables, we used both climatic and soil data. Climate data consisted of 19 bioclimatic variables related to temperature, rainfall and seasonality obtained from WorldClim 2.1 (https://www.worldclim.org; Fick & Hijmans, 2017), provided as gridded files at 2.5 arc-min resolution (ca. 20 km²) (Fig.1). These data encompass the period from 1970 to 2000 and were derived from interpolated information from 9,000–60,000 weather stations and satellite images (Fick & Hijmans, 2017). For plants only, we also included soil variables. Soil data (upper 0–30 cm depth range), such as pH and % sand, were derived from SoilGrids v2.0 (Poggio et al., 2021). In the “Ecological Niche Models” section in the Supplementary Material text and Supplementary Table S1, we detail the variable selection procedure and the specific variables used for each species’ ENM.

To model the distribution of each species, we used ensembles of ENMs fitted with species occurrences and environmental variables in the R package ‘sdm’, version 1.1.8 (Fig.1) (Naimi & Araújo, 2016). For ENM fitting, we used a combination of machine learning algorithms, including boosted regression trees (BRT; Friedman, 2001), maxLike (Royle et al., 2012), and random forests (RF; Breiman, 2001). To assess model performance, we used 75% of the occurrence records for calibration and 25% for model evaluation. Although our main study area and subsequent network analyses were restricted to the Cerrado of São Paulo State, we obtained georeferenced occurrences and calibrated and projected the ENMs across the entire extent of South America, because many of the species evaluated here occur not only in the Cerrado, but also across other South American biomes. This broader modelling extent helps to mitigate the problem of niche truncation, which can lead to an incomplete representation of species’ ecological niches by underestimating the range of environmental conditions under which species can occur, thereby reducing ENM transferability to future climates (Owens et al., 2013; Pearson & Dawson, 2003; Thuiller, 2024). Details on ENM building and parametrization are provided in the “Ecological Niche Models” section in the Supplementary Material text.

### 2.3 Species distribution forecasts

To assess how plants and their frugivores may rearrange their distributions in the future, we forecasted ensemble models under future climate change scenarios for the 2050s (2041–2060) and 2070s (2061–2080). Future climate projections were based on the Sixth Assessment Report of the Intergovernmental Panel on Climate Change (IPCC AR6; https://www.ipcc-data.org). We obtained these data from WorldClim 2.1 (https://www.worldclim.org; Fick & Hijmans, 2017) as gridded files at 2.5 arc-min resolution for two concentration pathways (SSPs), representing alternative trajectories of greenhouse gas emissions combined with socioeconomic contexts (IPCC, 2023). The first one is the SSP2-4.5 concentration pathway, hereafter referred to as the mitigation scenario, which refers to a scenario in which emission growth stabilizes by mid-century due to moderate climate policies, leading to a projected temperature rise of ca. 2.7 °C by the end of the century (IPCC, 2023). We also chose the SSP5-8.5, which is considered a “pessimistic” scenario, hereafter referred to as the business-as-usual (B.A.U.) scenario, that assumes fossil-fuel-driven development with continuously rising emissions, leading to ca. 4.4 °C higher temperatures by 2100 (IPCC, 2023). For each SSP, the IPCC report provides different climate models with distinct input parameters and methods. We selected five of these models (ACCESS-CM2, CMCC-ESM2, EC-EARTH3, MIROC6, and MRI-ESM2-0), chosen to capture the uncertainty in future climate projections for temperature and precipitation over South America (Almazroui et al., 2021). In the forecasting models, edaphic variables for plants were kept constant because no future soil scenarios were available. The outcomes of the models were binarized maps with the potential distribution of each species under current conditions, as well as under the mitigation and business-as-usual climate change scenarios (Fig.1). Details on the binarization procedure are provided in the “Ecological Niche Models” section in the Supplementary Material text.

### 2.3 Species range size change and displacement

The binarized ENM maps allowed us to forecast how climate change may affect species distributions (Fig.1). To quantify variation in range size within our study area, we compared the sum of each occupied cell areas (ca. 20 km²) between the present and each future climate scenario. Additionally, for each species and scenario, we calculated the geographic centroid of potential distribution within our study area, using the center of minimum distance (CMD). The CMD identifies the point among occupied cells that minimizes the total geodesic distance to all other occupied cells, making it particularly suitable for species with irregular or fragmented distributions. This analysis was performed using the “terra” R package, version 1.8.21 (Hijmans, 2025) and the “sf” R package, version 1.0.12 (Pebesma, 2018). To calculate species range displacement, i.e., the distance (km) between the current and future centroids, we measured distances on the surface of an ellipsoid (Vincenty, 1975). This calculation was performed using the “geosphere” R package, version 1.8 (Hijmans, 2022). The angular direction of centroid shifts was calculated using the bearing function from the “geosphere” R package, version 1.8 (Hijmans, 2022). This function computes the initial geodesic bearing between the current and future centroids on the Earth’s ellipsoid, yielding an angle in degrees relative to geographic north (0° = N, 90° = E, 180° = S, 270° = W). These bearings were standardized to the 0–360° range and subsequently transformed into compass directions: angles between 337.5° and 22.5° were classified as north (N), while angles from 22.5° to 67.5° were categorized as northeast (NE). East (E) included angles from 67.5° to 112.5°, and southeast (SE) covered angles from 112.5° to 157.5°. South (S) was defined for angles between 157.5° and 202.5°, and Southwest (SW) for angles from 202.5° to 247.5°. West (W) encompassed angles from 247.5° to 292.5°, and Northwest (NW) included angles from 292.5° to 337.5°.

In addition, we estimated the degree of species co-occurrence under the different climate change scenarios. To this end, we overlayed binary distribution maps for each plant-frugivore species pair under each climatic scenario and estimated their co-occurrence where both species were present (i.e., grid-cells with shared environmental suitability). These projections were used both to estimate potential pairwise species co-occurrence across the whole study area and to define the composition of local ecological communities

### 2.4 Species interaction probability matrix

To project plant-frugivore interactions under future climate scenarios and assemble local ecological networks, we first built an interaction probability matrix for the empirical species pool. This matrix combined three ecological mechanisms known to shape interaction frequency: temporal overlap, trait matching, and geographic range overlap (Vázquez et al., 2009; Vizentin-Bugoni & Maruyama, 2023). We then compared the predicted probabilities with empirical plant–frugivore interaction frequency data.

The s*pecies temporal overlap (T)* was estimated as the number of months each plant-frugivore species pair was simultaneously recorded, based on the fruit and frugivore phenology data (see *Study area and dataset* for details on phenology estimation). Hence, interaction probabilities increase with increasing plant-frugivore temporal overlap. To quantify *trait matching (M)*, we used fruit diameter, gape size of birds and gullet size of mammals, which are traits known to influence seed dispersal interactions (Bender et al., 2018). We used a modified Gower similarity index (Legendre & Legendre, 2012), in which the probability of interaction between frugivore *i* and plant *j* (P_ij_) is calculated as 1 minus the absolute difference between frugivore gape or gullet size (x_i_) and fruit diameter (y_i_), standardized by the range of trait values across all species in the network (*x* and *y*, Vizentin-Bugoni et al., 2019). Formally:

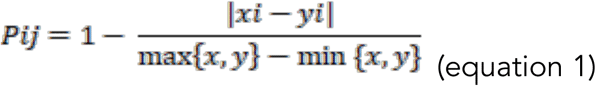

Hence, plant-frugivore partners with similar trait sizes are more likely to interact and the probability of interactions decreases as differences in partners’ traits increase. *Species range overlap (R)* was considered to be a proxy of the probability of plant-frugivore encounter in a local (grid-cell). Using our present-day binary maps, we calculated for each pairwise plant-frugivore combination the proportion of each plant species’ area that was co-occupied by each frugivore species. Thus, for the pair (animal *j*, plant *i*), we had:

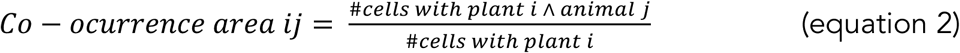

Finally, to give equal weight to the three matrices (*T*, *M*, *R*), we normalized each component matrix by its total sum (i.e., divided every entry by the sum of all entries in that matrix). Then, to create our interaction probability matrix, we multiplied the matrices element-wise (i.e., *T*\**M*\**R*) and normalized the resulting product (Vázquez et al., 2009; Vizentin-Bugoni et al., 2014). To assess whether these interaction probabilities reflected field-observed interaction frequencies, we compared our predicted interaction probability matrix with the empirical plant-frugivore interaction frequency matrix (see “Sampling of plant-frugivore interactions from the compiled dataset” in the Supplementary Material for details on how interaction frequency was obtained) (Campagnoli et al., 2025) - with a Mantel test. The predicted interaction probability matrix showed a strong correspondence with the empirical interaction frequency matrix (Mantel r = 0.74, p = 0.001, with 999 permutations), suggesting our model represented a fair approximation of the plant-frugivore interaction patterns observed.

To evaluate the influence of different climate-change scenarios on species interaction probabilities, we generated scenario-specific predictions. To this end, we replaced the species range overlap matrix, derived from animal-plant distributions in the binary maps and co-occurrence as in Eq. 2, with the one projected for each scenario while keeping the other two components (temporal overlap and trait matching) unchanged.

### 2.5 Grid-cell level plant-frugivore co-occurrence and local network assembly

ENMs were projected at ca. 21 km² resolution. To assemble local networks at a spatial grain comparable to the empirical interaction network sampled in Itirapina (ca. 50 km²), we aggregated binary distribution maps to cells of approximately 50 km². Each 50-km² grid cell was then treated as a local community, and local weighted networks were assembled by retaining only the plant and frugivore species predicted to co-occur within that cell. Link weights were assigned using the scenario-specific interaction probability matrix described above (Fig. 1), such that interaction strength for each species pair was given by the corresponding value in the global matrix (T × M × R, normalized). Because range overlap (R) varied across scenarios, interaction probabilities changed accordingly. Each cell on the map thus yielded an animal × plant matrix representing a spatially constrained subset of the full interaction matrix. We calculated network metrics only for cells containing at least two plant and two frugivore species (1,244 ± 83 cells; 92 ± 6% of all Cerrado–São Paulo cells per scenario), resulting in 6,221 local networks. Cells not meeting this criterion were excluded.

For each retained cell, we recorded the number of plant and frugivore species and the number of realized pairwise interactions, and two structural network descriptors. The weighted nestedness (*wNODF*) that indicates the extent to which interactions are organized around a generalist core, with specialist species interacting with a subset of partners generalists interact with, weighted by interaction probability (higher-probability links contribute more) (Almeida-Neto & Ulrich, 2011); and specialization (H2′) measures how much the observed interaction pattern deviates from expectations given partner availability, where availability is estimated as the sum of rows or columns of the observed matrix (Blüthgen et al., 2006). In our case, higher specialization indicates more unique (selective) interaction profiles and a more specialized network (Fig.1h).

### 2.6 Statistical analysis

To assess how climate change influences species’ range size and range displacement, we fitted four linear mixed models (LMMs). In the first set of models, we used plant and frugivore species range size, separately, as the response variable, while for the second set of models, we used the plant and frugivore range displacement (i.e., the distance in km between the current and future centroids of species distribution), respectively. Climate scenarios were the fixed predictor (with five levels: current, mitigation for the 2050s and 2070s, and business-as-usual for the 2050s and 2070s) in all models. These models included random intercepts for species to account for repeated measures across scenarios. Because exploratory diagnostics indicated heteroscedasticity among climate scenarios, we allowed residual variance to differ across scenarios using a variance structure (varIdent) implemented in the *nlme* package version 3.1 (Pinheiro & Bates, 1999). Models including heterogeneous variance structures were compared to homoscedastic models using AIC and likelihood ratio tests. In all cases, models allowing heterogeneous residual variance among climate scenarios were strongly supported (ΔAIC > 19; LRT p < 0.001) and were therefore retained for inference.

To evaluate whether changes in pairwise plant-frugivore co-occurrence were driven by changes in frugivore and plant range size and/or range shift, we fitted a LMM. We used the percentage of change in pairwise co-occurrence as the response variable, and the percent change in plant and frugivore range size and range displacement (centroid shift in km) as predictor variables. We included random intercepts for climate scenarios, plant, and frugivore species, as the model was fit on pairwise data (one record per plant-frugivore species pair x scenario).

Additionally, to test the effects of climate scenarios on interaction networks, we fitted a generalized linear mixed model (GLMM) for each network metric, with climate scenario as a fixed effect and random intercepts for cell identity (1|cell) to account for repeated measures across scenarios (Fig. 1). Because we detected spatial autocorrelation among neighboring cells (Moran’s *I*, *P* < 0.05), we included a spatial random field with a Matérn covariance using the spaMM R package, version 4.4.16 (Rousset et al., 2013), specified as Matern (1|x + y), where *x* and *y* are centered grid-cell centroid coordinates. For count responses (number of frugivores, plants, and links), we initially fitted models with a Poisson error distribution and tested for overdispersion using the *DHARMa* package, version 0.4.7 (Hartig et al., 2024). Because these models showed overdispersion, the final analyses for count responses were conducted using a negative binomial error distribution with a log link. Continuous response variables (weighted nestedness, and specialization) were analyzed with Gaussian error distributions after variance-stabilizing transformations: weighted nestedness was log-transformed, whereas specialization was square-root transformed. All LMM and GLMM models were performed using the “lme4” R package, version 1.1.34 (Bates et al., 2015).

## 3. RESULTS

Our models indicated a consistent and significant reduction in climatically suitable areas for plants and their seed dispersers under climate change scenarios in the Cerrado (within São Paulo state) compared to their current ranges (Table 1). On average, plants are expected to lose 62 ± 21% (mean ± SD) of their range under the mitigation scenario and 65 ± 26% under the business-as-usual scenario by the 2070s (Fig. 2a and Supplementary Table S2). Most plant species showed some level of contraction in distribution, while species such as *Davilla elliptica* and *Guapira noxia* are forecasted to lose nearly all suitable distribution under some climate change scenarios. For frugivores, by the 2070s we project losses of 32 ± 31% and 48 ± 36% under mitigation and business-as-usual scenarios, respectively (Fig. 2a and Supplementary Table S2). Some bird species, such as *Elaenia obscura* and *Hylophilus amaurocephalus*, and mammals such as Sus *scrofa*, are projected to become regionally extinct, with almost no climatic analogues in the future (Supplementary Table S2). Only two bird species (*Cnemotriccus fuscatus* and *Conirostrum speciosum*) showed no reduction in projected distribution, while a few species exhibited slight increases.

**Figure 2:**
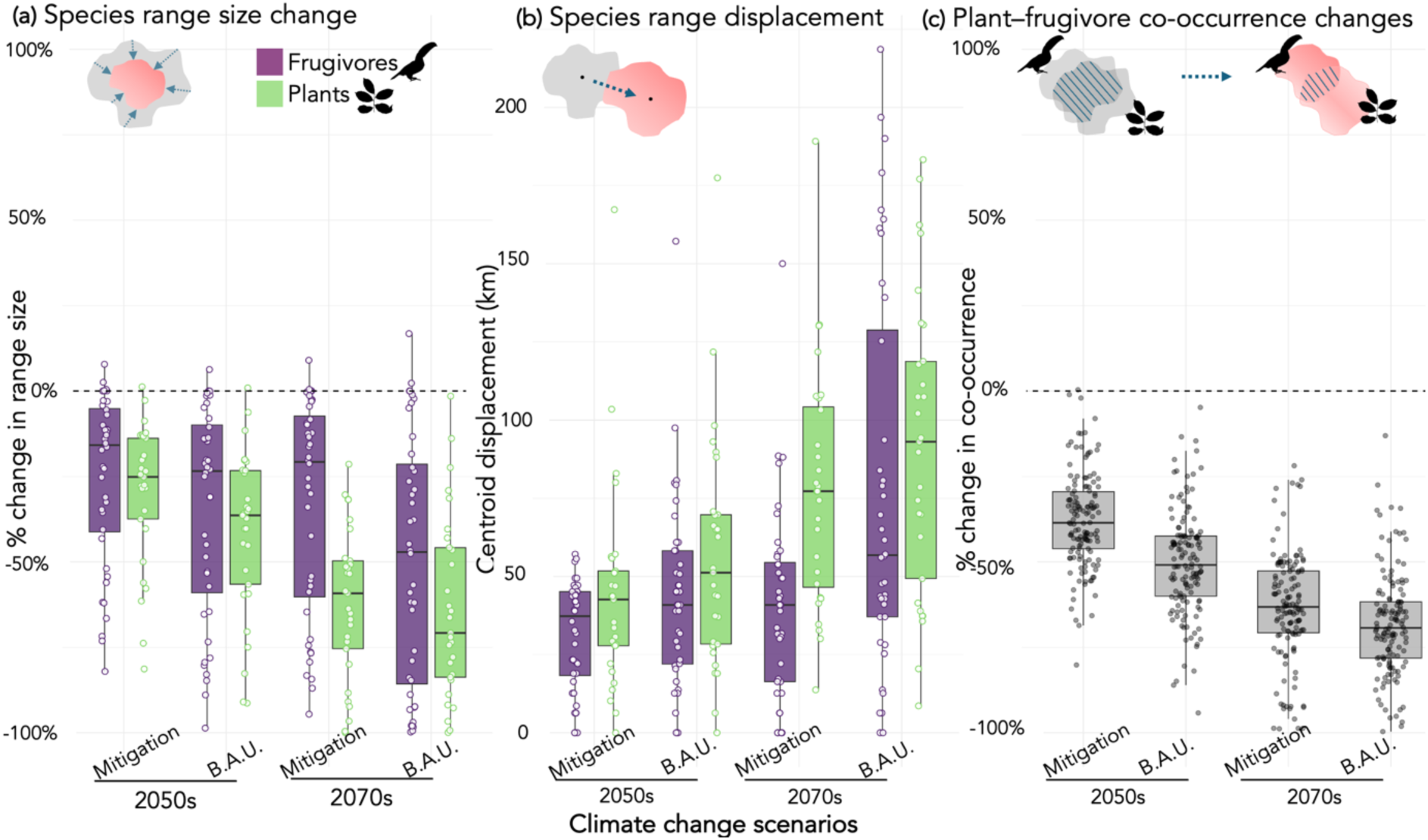
Climate change effects on species range size, displacement, and co-occurrence under mitigation and business-as-usual scenarios for the 2050s and 2070s compared to current scenario (baseline = 0). (a) Percentage changes in range size of frugivores (purple) and plants (green). (b) Species range displacement, measured as the distance (km) between the current and future centroids for frugivores and plants. (c) Percentage changes in plant–frugivore co-occurrence (i.e., proportion of plant range overlapping with their dispersers).

**Table 1:**
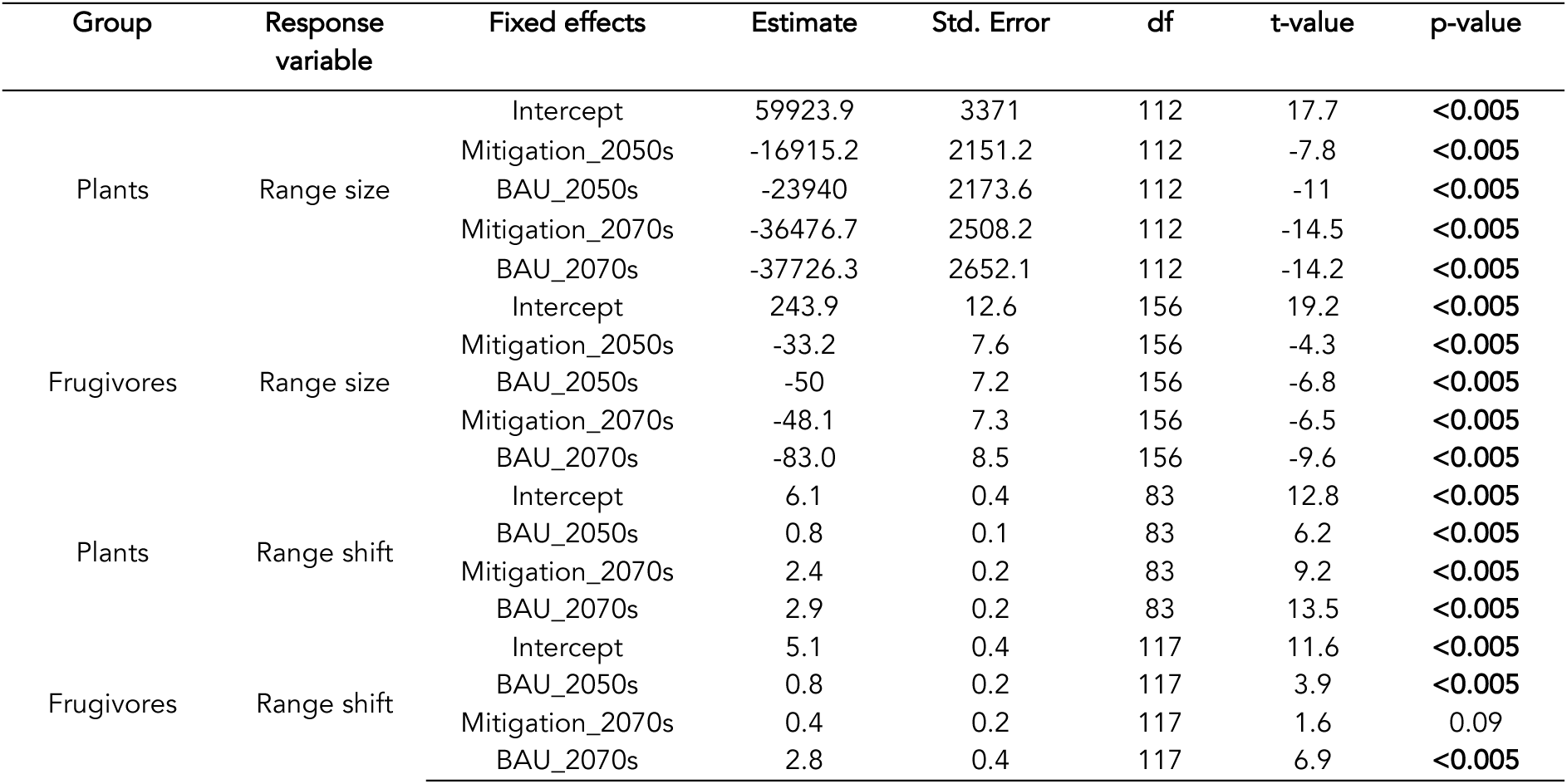
Future climate scenarios reduce species’ range size and increase range displacement. Coefficient estimates from linear mixed-effects models testing the effects of future climate scenarios on plant and frugivore range size and range displacement. Models included random intercepts for species and allowed heterogeneous residual variance across climate scenarios using a varIdent structure. Frugivore range size and both plant and frugivore range displacement were square root transformed, whereas plant range size was analysed on the original scale. Degrees of freedom correspond to denominator degrees of freedom estimated by the nlme R package (Pinheiro et al., 1999). For range-size models, the intercept corresponds to the current scenario. For range-displacement models, the intercept corresponds to the mitigation scenario for the 2050s, as current displacement is zero by definition. Bold values indicate statistically significant effects at **α** = 0.05.

**Table 2:** Drivers of changes in pairwise plant–frugivore co-occurrence. Coefficient estimates from a linear mixed model testing whether percentage changes in pairwise plant–frugivore co-occurrence were associated with plant and frugivore range-size change and range displacement. Range-size change was defined as the relative change in species’ range size, and range displacement as the distance between current and future range centroids. Plant species, frugivore species and scenario were included as random effects. Bold values indicate statistically significant effects at **α** = 0.05.

| Fixed effects | Estimate | Std. Error | df | Test value | p |
| --- | --- | --- | --- | --- | --- |
| Intercept | -19.65 | 2.70 | 14.64 | -7.26 | **< 0.001** |
| Plant range size change | 0.68 | 0.04 | 255.06 | 17.10 | **< 0.001** |
| Frugivorous range size change | 0.43 | 0.03 | 108.44 | 14.19 | **< 0.001** |
| Plant range displacement | 0.39 | 0.02 | 241.76 | 1.59 | 0.2 |
| Frugivorous range displacement | 0.06 | 0.01 | 505.19 | 3.32 | **< 0.001** |

We also found that species not only contract their distributions but also undergo a significant range shift under future scenarios (Table 1), with shifts in the 2070s being the most pronounced (Fig. 2b and Supplementary Table S3). Specifically, frugivore range centroids are forecasted to shift by 41 ± 31 km and 77 ± 61 km under mitigation and business-as-usual scenarios, respectively, by the 2070s compared to their current locations. Meanwhile, plant centroid shifts are predicted to be even larger, with 78 ± 40 km range displacement under the mitigation scenario and 91 ± 49 km under the business-as-usual scenario (Fig. 2b and Supplementary Table S3). The majority of plants (85–90%) and their seed dispersers (60–70%) are projected to experience centroid shifts toward the southern and southwestern portions of the Cerrado, although for some species we observed multidirectional range shifts toward multiple directions (Supplementary Table S3).

Our forecast also suggests a substantial shrinkage in plant-frugivore co-occurrence (Fig. 2c and Supplementary Table S4). Under the mitigation scenario by the 2070s, 19 out of the 139 pairwise interactions are projected to lose 25-50% of their co-occurrence area, 70 to lose 50-75%, 48 to lose 75-100%, and only 2 to remain stable or lose up to 25%. However, under a continuously rising greenhouse gas emissions scenario, represented by the business-as-usual scenario, 9 out of the 139 plant-frugivore interactions are forecasted to lose 25-50% of their co-occurrence area, 46 to lose 50-75%, and 83 to lose 75-100%. In all scenarios, no pairwise interaction is expected to increase its co-occurrence compared to current conditions (Fig. 2c and Supplementary Table S4). Changes in both plant and frugivore range size significantly explained variation in pairwise co-occurrence (plant range size: *t* = 17.10, *p* < 0.001; frugivore range size: *t* = 14.20, *p* < 0.001), with plant range shifts showing the strongest effect.

These climate-driven changes in species co-occurrence translated into significantly smaller local Cerrado plant–frugivore networks (Table 3; Figure 3). For instance, relative to the current scenario, the mean number of frugivores per local network declined by 59%, from 34 ± 6 to 14 ± 7 species, under the business-as-usual scenario for the 2070s. Similarly, the mean number of plant species declined by 65%, from 26 ± 4 to 9 ± 6 species under the same scenario. The number of links was also significantly lower in all future scenarios compared with present-day networks. In addition to these reductions in network size, weighted nestedness declined significantly across all future scenarios, with increasingly negative effects under later and more severe climate projections (Table 3). In contrast, specialization increased significantly in all future scenarios, indicating that future local networks are expected to become more specialized than current networks (Table 3). Overall, these results suggests that climate change is expected to simplify the structure of local Cerrado plant–frugivore networks, reducing their size, number of interactions, and hierarchical organization, while increasing specialization (Figure 3).

**Figure 3:**
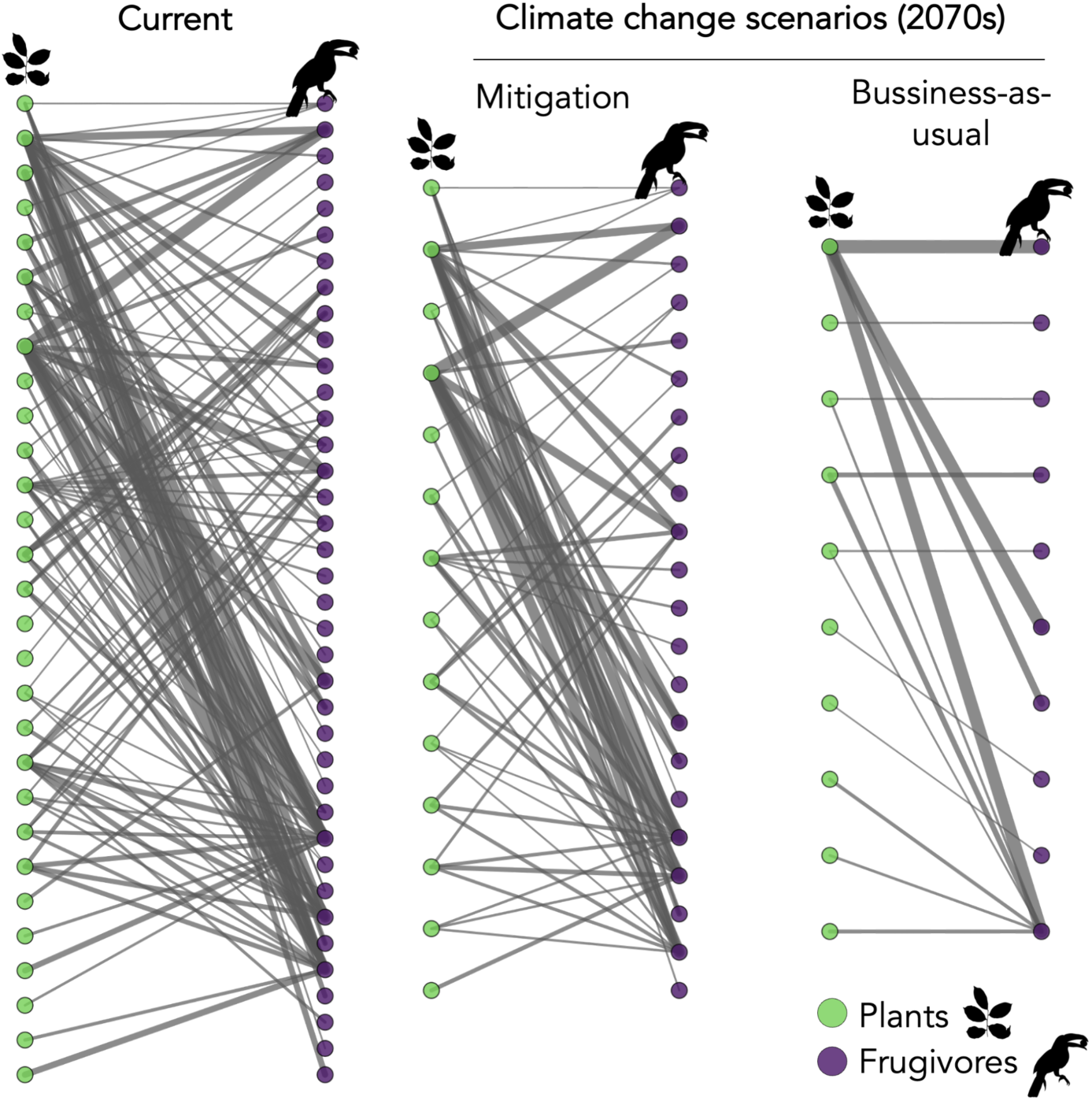
Climate change simplifies and reduces local Cerrado plant–frugivore networks. Illustration depicts one representative local network from the Cerrado under current conditions and future climate scenarios for the 2070s. The same focal cell is displayed across scenarios to illustrate how climate-driven changes in species co-occurrence translate into smaller and structurally simpler local interaction networks under mitigation and business-as-usual projections, with a reduction in the number of links and species richness. Green nodes represent plants, purple nodes represent frugivorous birds, and link width is proportional to interaction probability.

**Table 3:** Effects of future climate scenarios on local network structure. Coefficient estimates from spatial generalized linear mixed models testing the effects of future climate scenarios on frugivore richness, plant richness, number of links, weighted nestedness (wNODF), and specialization (H₂′) in local Cerrado plant-frugivore networks. The current scenario was used as the reference level. Models included cell identity as a random effect and a spatial random field with a Matérn covariance structure. Bold values indicate statistically significant effects at **α** = 0.05.

| Network metrics | Fixed effects | Estimate | Std. Error | Z-value | p-value |
| --- | --- | --- | --- | --- | --- |
| Number of frugivorous species | Intercept | 3.29 | 0.11 | 28.6 | **<0.005** |
|  | Mitigation_2050s | -0.33 | 0.007 | -42.8 | **<0.005** |
|  | Mitigation_2070s | -0.61 | 0.008 | -70.6 | **<0.005** |
|  | BAU_2050s | -0.51 | 0.008 | -61.9 | **<0.005** |
|  | BAU_2070s | -0.93 | 0.009 | -98.4 | **<0.005** |
| Number of plant species | Intercept | 3.05 | 0.11 | 27.6 | **<0.005** |
|  | Mitigation_2050s | -0.39 | 0.01 | -35.6 | **<0.005** |
|  | Mitigation_2070s | -0.94 | 0.01 | -73.8 | **<0.005** |
|  | BAU_2050s | -0.59 | 0.01 | -51.6 | **<0.005** |
|  | BAU_2070s | -1.11 | 0.01 | -85.0 | **<0.005** |
| Number of links | Intercept | 4.54 | 0.165 | 27.6 | **<0.005** |
|  | Mitigation_2050s | -0.621 | 0.015 | -39.7 | **<0.005** |
|  | Mitigation_2070s | -1.18 | 0.016 | -69.6 | **<0.005** |
|  | BAU_2050s | -0.884 | 0.015 | -55.4 | **<0.005** |
|  | BAU_2070s | -1.53 | 0.016 | -91.2 | **<0.005** |
| Weighted nestedness | Intercept | 2.90 | 0.30 | 9.53 | **<0.005** |
|  | Mitigation_2050s | -0.27 | 0.05 | -5.19 | **<0.005** |
|  | Mitigation_2070s | -0.41 | 0.05 | -7.37 | **<0.005** |
|  | BAU_2050s | -0.29 | 0.05 | -5.43 | **<0.005** |
|  | BAU_2070s | -1.08 | 0.05 | -19.6 | **<0.005** |
| Specialization ($H_2'$) | Intercept | 0.60 | 0.016 | 36.9 | **<0.005** |
|  | Mitigation_2050s | 0.018 | 0.0031 | 5.78 | **<0.005** |
|  | Mitigation_2070s | 0.042 | 0.0032 | 12.7 | **<0.005** |
|  | BAU_2050s | 0.020 | 0.0031 | 6.54 | **<0.005** |
|  | BAU_2070s | 0.083 | 0.0032 | 26.0 | **<0.005** |

## 4. DISCUSSION

Long-term changes in climate patterns, primarily driven by human activities in the last century, are impacting biodiversity and ecosystems. Our study demonstrates that future climate may pose a threat not only to species distributions but also to ecological interactions, such as seed dispersal, that are key to enabling plants to track suitable climate. Thus, preventing the simplification of interaction networks will be essential to preserve biodiversity in species-rich regions like the Cerrado.

Future climatic projections indicate that the Cerrado may experience an extension of the dry season, with some regions facing up to a 15% reduction in rainfall, accompanied by increasing temperatures (Hofmann et al., 2025). Under such a hotter and drier future, the geographic distributions of both plants and animals that compose our empirical seed-dispersal network are expected to contract and shift substantially. These findings corroborate previous studies that project future range contractions and biotic homogenization among Cerrado communities of birds (Hidasi-Neto et al., 2022), mammals (Hidasi-Neto et al., 2019) and plants (Velazco et al., 2019). While mammal homogenization and severe bird species loss are particularly expected in the southern portion of the Cerrado, our focal study region, bird homogenization is also projected to occur further north (Hidasi-Neto et al., 2019). These projections are particularly alarming because much of the Cerrado’s biodiversity is already threatened by the conversion of natural habitats to agriculture and exotic pasturelands, as well as increases in anthropogenic wildfires, making the biome one of the most vulnerable tropical ecosystems (Hofmann et al., 2025; Strassburg et al., 2017). These anthropogenic pressures likely further reduce landscape permeability. Because projected climate change is likely to outpace in situ adaptive responses for most species (Quintero & Wiens, 2013; Radchuk et al., 2019), many populations may become increasingly confined to suboptimal environments, potentially leading to population declines and, ultimately, local extinctions. In this context, climate change is likely to exacerbate the threats already faced by Cerrado biodiversity.

Co-occurring animals and plants do not necessarily respond to global change in the same way. Climate change can thus disrupt trophic interactions through the decoupling of the geographic distributions of interacting partners (Sales, Galetti, et al., 2020). Here, we project a severe loss of co-occurrence between plants and their dispersers across the Cerrado. For example, under the most severe scenario for the 2070s, we estimated that 60% of the interactions (83 out of 138) would be severely threatened, with the interaction partners losing 75–100% of their co-occurrence area. The decline of frugivores can lead to substantial consequences for plant demography by lowering plant recruitment, decreasing local plant diversity (Gardner et al., 2019), and causing greater spatial clustering of plant species across landscapes, which can increase density-dependent mortality caused by herbivores or pathogens (Carvalho et al., 2021; Fricke & Wright, 2017).

Because plants are sessile organisms, animals are critical in helping them expand their distributions toward more favorable conditions (Corlett & Westcott, 2013). Our models forecasted, for example, that some plant species might need to shift their distributions by more than 100 km over the coming decades, which would be impossible to achieve naturally without animal-assisted dispersal. Some birds that perform migratory movements are important seed dispersers in Cerrado, including migrants coming from Northern (e.g., *Vireo olivaceus*, *Elaenia spectabilis*) and Southern locations (e.g., *Turdus amaurochalinus*). These birds may be especially important to allow plants to colonize suitable sites displacing their ranges north-south to follow their suitable climate niches (González-Varo et al., 2021). In addition to the direction of seed dispersal, reductions in dispersal distances can seriously compromise the ability of plants to respond to global change by shifting their ranges (Nuñez et al., 2023). A decrease in seed dispersal distances may be exacerbated by defaunation (i.e., the loss of individuals, populations, and species of animals), which has globally reduced the plant’s capacity to track climate by 60% (Fricke et al., 2022). Defaunation may not only reduce the distances of dispersal achieved, but also the number of seeds transported away and the quality of dispersal (see (Campagnoli, Rumeu, et al., 2025; Jacques et al., 2025 for examples in Cerrado). The severe reduction in co-occurrence projected in our study may therefore create a vulnerability loop, in which plant responses to climate change become increasingly constrained by both defaunation and the loss of spatial overlap with seed dispersers.

For animals, the shrinkage of co-occurrence with plants may represent an important nutritional constraint. For instance, fruits can constitute more than half of the diet of some frugivores, including *Antilophia galeata* (Marini, 1992), *Penelope superciliaris* (Zaca et al., 2006)*, Elaenia cristata (Guerra & Pizo, 2014)*, *Turdus amaurochalinus* and *Turdus leucomelas (Gasperin & Aurélio Pizo, 2009)*. Even for some omnivorous dispersers from our Cerrado seed dispersal network, fruits may act as a critical resource during the dry season, e.g., for *Didelphis albiventris (Cantor et al., 2010)*. Together, these results suggest that climate change may erode not only species distributions, but also the ecological interactions that sustain plant persistence and animal nutrition.

At the community level, we forecast that climate-driven reductions in species co-occurrence may alter the composition of ecological assemblages and reshape plant-animal interactions across the Cerrado. Under future climate scenarios, local communities are expected to contain fewer plant and frugivore species, as well as fewer interaction links, indicating a marked contraction of local ecological networks. These results are consistent with previous studies showing that environmental disturbances such as fragmentation, habitat loss, and logging can reduce species richness and interaction diversity in seed-dispersal systems (e.g.,Emer et al., 2020; Li et al., 2022; Rossi et al., 2025). Our results indicate another layer of perturbation on Cerrado networks. Indeed, we found that climate change tends to erode the architecture of local mutualistic networks, with effects becoming progressively stronger under more severe future scenarios. Specifically, future scenarios led to significant declines in weighted nestedness, indicating that networks become less hierarchically organized. This reduction in nestedness suggests that specialist species no longer interact with subsets of generalist partners, weakening the cohesive structure that usually characterizes mutualistic networks (Almeida-Neto et al., 2008; Bascompte et al., 2003). Concurrently, our models projected a significant increase in network specialization under future scenarios. In a less nested network, higher specialization suggests that the remaining interactions become more unevenly partitioned among species, with reduced overlap in partner use (Bascompte et al., 2003; Blüthgen et al., 2006; Schleuning et al., 2020; Vázquez & Aizen, 2004). Thus, because highly nested networks are expected to promote stability in mutualistic communities, the simultaneous decline in nestedness and increase in specialization suggest that future Cerrado networks may be less stable and resilient than present-day ones (Thébault & Fontaine, 2010). Together, these results suggest that climate change will not only reduce the size of Cerrado plant-frugivore networks but also simplify their organization in ways that may cause lower stability and resilience.

It is important to note that our projections constrained potential interactions to those in our empirical network, which was limited by the species pool and interactions detected in the field (although our high sampling effort was substantial and included a combination of different sampling methods). Therefore, projected losses and the simplification of Cerrado local networks may be partially buffered if species are highly capable of switching to alternative resources after losing a primary partner, a process known as interaction rewiring (Poisot et al., 2012; Vizentin-Bugoni & Maruyama, 2023). However, spatial co-occurrence of potential partners is a critical precondition for an interaction - and, therefore, rewiring - to occur (Jordano, 2016). Although rewiring is possible, its capacity to compensate losses in seed dispersal systems still requires empirical testing (Vizentin-Bugoni & Maruyama, 2023, but see Costa et al., 2018; Timóteo et al., 2016), as simulations often fail to incorporate species’ ecological plasticity (such as omnivory), complex behavioural and population-level responses, and whether newly formed interactions effectively preserve ecological function (Costa et al., 2018). Thus, even if interaction rewiring allows network structure to remain relatively stable, there is no guarantee that ecological functions will be retained (Timóteo et al., 2016; Vizentin-Bugoni & Maruyama, 2023). Additionally, our interaction probability matrix assumed that temporal overlap between plants and frugivores (i.e., phenological overlap) remained constant across climate scenarios. However, recent evidence indicates that climate change may delay or advance fruiting periods and substantially alter the duration of fruiting (Ji et al., 2026; Martins et al., 2025a). These phenological changes imply that temporal overlap between interacting partners may also change across future climate scenarios, potentially disrupting the temporal overlap between plants and their seed dispersers (Sandor et al., 2021). Although not accounting for this may constrain our predictions, there is large interspecific variability in phenological responses to climate shifts (Ji et al., 2026). Forecasting these complex phenological changes in highly diverse tropical systems remains a major challenge, and there is currently insufficient long-term data to accurately incorporate these variables into our community-level modeling (Martins et al., 2025b; Sandor et al., 2021).

By integrating empirical data on plant-frugivore interaction data with species distribution models, we show that climate change may deeply reorganize local interaction networks in the world’s most diverse savanna. Projected networks are smaller, structurally simpler, and less resilient, revealing that the impacts of climate change may extend beyond species range shifts to the erosion of ecological functioning itself. Because seed dispersal underpins natural regeneration (Estrada-Villegas et al., 2023), carbon storage (Bello et al., 2015), and the capacity of plants to track suitable climates (Nuñez et al., 2023), its disruption may compromise both biodiversity persistence and ecosystem resilience. Our results therefore highlight that climate change threatens not only species, but also the ecological interactions that sustain ecosystem dynamics and long-term conservation outcomes.

## Supporting information

Supplementary informations

## Acknowledgements

This research was partially supported by the Coordenação de Aperfeiçoamento de Pessoal de Nível Superior - Brasil (CAPES), through Finance Code 001 and the CAPES PrInt program (grant #88882.426377/2019-01). We gratefully acknowledge the Rufford Foundation (grant no. 33432-1) and the Neotropical Grasslands Conservancy for their financial support toward equipment acquisition and field activities. GP was funded by Agência Nacional de Promoción de la Investigación, el Desarrollo Tecnológico y la Innovación (#PICT-2021-I-INVI-00071) and JVB by Instituto Serrapilheira (Serra 2211-42118) and FAPERGS (T.O. 23/2551-0001592-7). AVC is currently supported by Fundação de Amparo à Pesquisa do Estado de São Paulo (FAPESP #2023/16620-0) and Conselho Nacional de Desenvolvimento Científico e Tecnológico (CNPq # 307439/2025-9). We are also thankful to the staff of the Estação Ecológica e Experimental de Itirapina for their assistance throughout the study, and to the Fundação Florestal for granting the necessary research permit.

## Contributions

E.D.B.R., M.C., J.V.B., and G.P. conceived the study. M.C and A.V.C collected the data, E.D.B.R. and M.C. performed the statistical analyses. E.D.B.R. led manuscript writing and data visualization, with equal contributions from M.C., J.V.B., A.V.C., and G.P. All authors contributed to manuscript revision and editing.

## Data availability statement

The climate data used in this word can be downloaded from CHELSA (https://chelsa-climate.org). The soil data can be downloaded from SOILGRIDS (https://soilgrids.org). Species interaction data, as well, as the R code used for species distribution modelling and linear models will be publicly available after peer-review.

## Data availability statement

The climatic and soil predictor rasters, as well as the curated species occurrence records used for the analyses, are available via Zenodo at xxxx.

The R code used for species occurrence acquisition, cleaning, and spatial thinning, as well as code for species distribution modelling, richness- and function-based predictions, and regression analyses, is available via Zenodo at https://doi.org/10.5281/zenodo.17054306

## Reference

Almazroui, M., Ashfaq, M., Islam, M. N., Rashid, I. U., Kamil, S., Abid, M. A., O’Brien, E., Ismail, M., Reboita, M. S., Sörensson, A. A., Arias, P. A., Alves, L. M., Tippett, M. K., Saeed, S., Haarsma, R., Doblas-Reyes, F. J., Saeed, F., Kucharski, F., Nadeem, I., … Sylla, M. B. (2021). Assessment of CMIP6 Performance and Projected Temperature and Precipitation Changes Over South America. Earth Systems and Environment, 5(2), 155–183. 10.1007/s41748-021-00233-6

Almeida-Neto, M., Guimarães, P., Guimarães, P. R., Loyola, R. D., & Ulrich, W. (2008). A consistent metric for nestedness analysis in ecological systems: reconciling concept and measurement. Oikos, 117(8), 1227–1239. 10.1111/j.0030-1299.2008.16644.x

Almeida-Neto, M., & Ulrich, W. (2011). A straightforward computational approach for measuring nestedness using quantitative matrices. Environmental Modelling & Software, 26(2), 173–178. 10.1016/j.envsoft.2010.08.003

Bascompte, J., Jordano, P., Melián, C. J., & Olesen, J. M. (2003). The nested assembly of plant–animal mutualistic networks. Proceedings of the National Academy of Sciences, 100(16), 9383–9387. 10.1073/pnas.1633576100

Bates, D., Mächler, M., Bolker, B., & Walker, S. (2015). Fitting Linear Mixed-Effects Models Using lme4. Journal of Statistical Software, 67(1). 10.18637/jss.v067.i01

Bello, C., Galetti, M., Montan, D., Pizo, M. A., Mariguela, T. C., Culot, L., Bufalo, F., Labecca, F., Pedrosa, F., Constantini, R., Emer, C., Silva, W. R., da Silva, F. R., Ovaskainen, O., & Jordano, P. (2017). Atlantic frugivory: a plant–frugivore interaction data set for the Atlantic Forest. Ecology, 98(6), 1729. 10.1002/ecy.1818

Bello, C., Galetti, M., Pizo, M. A., Magnago, L. F. S., Rocha, M. F., Lima, R. A. F., Peres, C. A., Ovaskainen, O., & Jordano, P. (2015). Defaunation affects carbon storage in tropical forests. Science Advances, 1(11). 10.1126/sciadv.1501105

Bender, I. M. A., Kissling, W. D., Blendinger, P. G., Böhning-Gaese, K., Hensen, I., Kühn, I., Muñoz, M. C., Neuschulz, E. L., Nowak, L., Quitián, M., Saavedra, F., Santillán, V., Töpfer, T., Wiegand, T., Dehling, D. M., & Schleuning, M. (2018). Morphological trait matching shapes plant–frugivore networks across the Andes. Ecography, 41(11), 1910–1919. 10.1111/ecog.03396

Blüthgen, N., Menzel, F., & Blüthgen, N. (2006). Measuring specialization in species interaction networks. BMC Ecology, 6. 10.1186/1472-6785-6-9

Breiman, L. (2001). Random Forests. Machine Learning, 45(1), 5–32. 10.1023/A:1010933404324

Calvin, K., Dasgupta, D., Krinner, G., Mukherji, A., Thorne, P. W., Trisos, C., Romero, J., Aldunce, P., Barrett, K., Blanco, G., Cheung, W. W. L., Connors, S., Denton, F., Diongue-Niang, A., Dodman, D., Garschagen, M., Geden, O., Hayward, B., Jones, C., … Ha, M. (2023). IPCC, 2023: Climate Change 2023: Synthesis Report. Contribution of Working Groups I, II and III to the Sixth Assessment Report of the Intergovernmental Panel on Climate Change [Core Writing Team, H. Lee and J. Romero (eds.)]. *IPCC*, Geneva, Switzerland. (P. Arias, M. Bustamante, I. Elgizouli, G. Flato, M. Howden, C. Méndez-Vallejo, J. J. Pereira, R. Pichs-Madruga, S. K. Rose, Y. Saheb, R. Sánchez Rodríguez, D. Ürge-Vorsatz, C. Xiao, N. Yassaa, J. Romero, J. Kim, E. F. Haites, Y. Jung, R. Stavins, … C. Péan, Eds.). Cambridge University Press. 10.59327/IPCC/AR6-9789291691647

Campagnoli, M., Christianini, A., & Peralta, G. (2025). Plant and frugivore species characteristics drive frugivore contributions to seed dispersal effectiveness in a hyperdiverse community. Functional Ecology, 39(1), 238–253. 10.1111/1365-2435.14697

Campagnoli, M., Rumeu, B., Peralta, G., & Christianini, A. (2025). Functional robustness declines faster than structural robustness in hyperdiverse seed dispersal networks following defaunation. Biological Conservation, 308, 111265. 10.1016/j.biocon.2025.111265

Cantor, M., Ferreira, L. A., Silva, W. R., & Setz, E. Z. F. (2010). Potential seed dispersal by Didelphis albiventris (Marsupialia, Didelphidae) in highly disturbed environment. Biota Neotropica, 10(2), 45–51. 10.1590/S1676-06032010000200004

Carvalho, C. da S., García, C., Lucas, M. S., Jordano, P., & Côrtes, M. C. (2021). Extant fruit-eating birds promote genetically diverse seed rain, but disperse to fewer sites in defaunated tropical forests. Journal of Ecology, 109(2), 1055–1067. 10.1111/1365-2745.13534

Comita, L. S., Queenborough, S. A., Murphy, S. J., Eck, J. L., Xu, K., Krishnadas, M., Beckman, N., & Zhu, Y. (2014). Testing predictions of the Janzen– Connell hypothesis: a meta-analysis of experimental evidence for distance- and density-dependent seed and seedling survival. Journal of Ecology, 102(4), 845–856. 10.1111/1365-2745.12232

Corlett, R. T., & Westcott, D. A. (2013). Will plant movements keep up with climate change? Trends in Ecology and Evolution, 28(8), 482–488. 10.1016/j.tree.2013.04.003

Costa, J. M., Ramos, J. A., da Silva, L. P., Timóteo, S., Andrade, P., Araújo, P. M., Carneiro, C., Correia, E., Cortez, P., Felgueiras, M., Godinho, C., Lopes, R. J., Matos, C., Norte, A. C., Pereira, P. F., Rosa, A., & Heleno, R. H. (2018). Rewiring of experimentally disturbed seed dispersal networks might lead to unexpected network configurations. Basic and Applied Ecology, 30, 11–22. 10.1016/j.baae.2018.05.011

Dehling, D. M., Töpfer, T., Schaefer, H. M., Jordano, P., Böhning-Gaese, K., & Schleuning, M. (2014). Functional relationships beyond species richness patterns: Trait matching in plant-bird mutualisms across scales. Global Ecology and Biogeography, 23(10), 1085–1093. 10.1111/geb.12193

Durigan, G., & Ratter, J. A. (2006). Successional changes in cerrado and cerrado/forest ecotonal vegetation in western São Paulo State, Brazil, 1962-2000. Edinburgh Journal of Botany, 63(1), 119–130. 10.1017/S0960428606000357

Emer, C., Jordano, P., Pizo, M. A., Ribeiro, M. C., da Silva, F. R., & Galetti, M. (2020). Seed dispersal networks in tropical forest fragments: Area effects, remnant species, and interaction diversity. Biotropica, 52(1), 81–89. 10.1111/btp.12738

Estrada-Villegas, S., Stevenson, P. R., López, O., DeWalt, S. J., Comita, L. S., & Dent, D. H. (2023). Animal seed dispersal recovery during passive restoration in a forested landscape. Philosophical Transactions of the Royal Society B: Biological Sciences, 378(1867). 10.1098/rstb.2021.0076

Fick, S. E., & Hijmans, R. J. (2017). WorldClim 2: new 1-km spatial resolution climate surfaces for global land areas. International Journal of Climatology, 37(12), 4302–4315. 10.1002/joc.5086

Filgueiras, B. K. C., Peres, C. A., Melo, F. P. L., Leal, I. R., & Tabarelli, M. (2021). Winner–Loser Species Replacements in Human-Modified Landscapes. Trends in Ecology & Evolution, 36(6), 545–555. 10.1016/j.tree.2021.02.006

Fricke, E. C., Ordonez, A., Rogers, H. S., & Svenning, J.-C. (2022). The effects of defaunation on plants’ capacity to track climate change. Science, 375(6577), 210–214. 10.1126/science.abk3510

Fricke, E. C., & Wright, S. J. (2017). Measuring the demographic impact of conspecific negative density dependence. Oecologia, 184(1), 259–266. 10.1007/s00442-017-3863-y

Friedman, J. H. (2001). Greedy function approximation: A gradient boosting machine. The Annals of Statistics, 29(5). 10.1214/aos/1013203451

Fuzessy, L., Sobral, G., Carreira, D., Rother, D. C., Barbosa, G., Landis, M., Galetti, M., Dallas, T., Cardoso Cláudio, V., Culot, L., & Jordano, P. (2022). Functional roles of frugivores and plants shape hyper-diverse mutualistic interactions under two antagonistic conservation scenarios. Biotropica, 54(2), 444–454. 10.1111/btp.13065

Gardner, C. J., Bicknell, J. E., Baldwin-Cantello, W., Struebig, M. J., & Davies, Z. G. (2019). Quantifying the impacts of defaunation on natural forest regeneration in a global meta-analysis. Nature Communications, 10(1), 1–7. 10.1038/s41467-019-12539-1

Gasperin, G., & Aurélio Pizo, M. (2009). Frugivory and habitat use by thrushes (Turdus spp.) in a suburban area in south Brazil. Urban Ecosystems, 12(4), 425–436. 10.1007/s11252-009-0090-2

Goel, N., Guttal, V., Levin, S. A., & Staver, A. C. (2020). Dispersal Increases the Resilience of Tropical Savanna and Forest Distributions. The American Naturalist, 195(5), 833–850. 10.1086/708270

Gonçalves, F., Sales, L. P., Galetti, M., & Pires, M. M. (2021). Combined impacts of climate and land use change and the future restructuring of Neotropical bat biodiversity. Perspectives in Ecology and Conservation, 19(4), 454–463. 10.1016/j.pecon.2021.07.005

González-Varo, J. P., Rumeu, B., Albrecht, J., Arroyo, J. M., Bueno, R. S., Burgos, T., da Silva, L. P., Escribano-Ávila, G., Farwig, N., García, D., Heleno, R. H., Illera, J. C., Jordano, P., Kurek, P., Simmons, B. I., Virgós, E., Sutherland, W. J., & Traveset, A. (2021). Limited potential for bird migration to disperse plants to cooler latitudes. Nature, 595(7865), 75–79. 10.1038/s41586-021-03665-2

Gottsberger, G., & Silberbauer-Gottsberger, I. (1983). Dispersal and distribution in the cerrado vegetation of Brazil. Sonderbänd Des Naturwissenschaftlichen Vereins in Hamburg, 7(2), 315–352.

Guerra, T. J., & Pizo, M. A. (2014). Asymmetrical Dependence Between a Neotropical Mistletoe and its Avian Seed Disperser. Biotropica, 46(3), 285–293. 10.1111/btp.12112

Guisan, A., & Zimmermann, N. E. (2000). Predictive habitat distribution models in ecology. Ecological Modelling, 135(2–3), 147–186. 10.1016/S0304-3800(00)00354-9

Hartig, F., Lohse, L., & de Souza leite, M. (2024). DHARMa: Residual Diagnostics for Hierarchical (Multi-Level / Mixed) Regression Models (0.4.7).

Hidasi-Neto, J., Gomes, N. M. A., & Pinto, N. S. (2022). Cerrado native vegetation is a refuge for birds under the current climate change trajectory. Austral Ecology, 47(8), 1622–1635. 10.1111/aec.13242

Hidasi-Neto, J., Joner, D. C., Resende, F., Monteiro, L. de M., Faleiro, F. V., Loyola, R. D., & Cianciaruso, M. V. (2019). Climate change will drive mammal species loss and biotic homogenization in the Cerrado Biodiversity Hotspot. Perspectives in Ecology and Conservation, 17(2), 57–63. 10.1016/j.pecon.2019.02.001

Hijmans, R. J. (2022). geosphere: Spherical Trigonometry. Https://CRAN.R-Project.Org/Package=geosphere.

Hofmann, G. S., Silva, R. C., Weber, E. J., Barbosa, A. A., Oliveira, L. F. B., Alves, R. J. V., Hasenack, H., Schossler, V., Aquino, F. E., & Cardoso, M. F. (2023). Changes in atmospheric circulation and evapotranspiration are reducing rainfall in the Brazilian Cerrado. Scientific Reports, 13(1), 11236. 10.1038/s41598-023-38174-x

Hofmann, G. S., Weber, E. J., Bastazini, V. A. G., Rossatto, D. R., Franco, A. C., Granada, C. E., Kaminski, L. A., Ubaid, F. K., Leandro-Silva, V., Borges-Martins, M., Silva, R. C., Cardoso, M. F., Oliveira, L. F. B., Aquino, F. E., & Pereira, M. J. R. (2025). Climate Change in the Brazilian Cerrado: A Looming Threat to Terrestrial Biodiversity. In Wiley Interdisciplinary Reviews: Climate Change (Vol. 16, Number 5). John Wiley and Sons Inc. 10.1002/wcc.70022

Howe, H. F., & Smallwood, J. (1982). Ecology of seed dispersal. Annual Review of Ecology and Systematics, 13(1), 201–228. 10.1146/annurev.es.13.110182.001221

Jacques, C. L. B., Mariano, V., & Christianini, A. V. (2025). Gut-passage time and the role of the Greater Rhea (Rhea americana) influencing seed dispersal and predation in Neotropical grasslands and savannas. Emu - Austral Ornithology, 125(1), 60–72. 10.1080/01584197.2024.2442343

Ji, G., Peng, Y., Li, X., Zhang, S., Zhu, H., Yang, Z., Lv, M., Berninger, F., & Ma, Q. (2026). Climate warming advances flowering and fruiting but drives divergent changes in reproductive season length. Communications Earth & Environment. 10.1038/s43247-026-03374-6

Jordano, P. (2000). Fruits and frugivory . In M. Fenner (Ed.), Seeds: the ecology of regeneration in plant communities (2nd ed., pp. 125–165). CABI Publishing. 10.1079/9780851994321.0125

Jordano, P. (2016). Sampling networks of ecological interactions. Functional Ecology, 30(12), 1883–1893. 10.1111/1365-2435.12763

Legendre, P., & Legendre, L. (2012). References (Vol. 24, pp. 907–968). 10.1016/B978-0-444-53868-0.50018-6

Li, W., Zhu, C., Grass, I., Vázquez, D. P., Wang, D., Zhao, Y., Zeng, D., Kang, Y., Ding, P., & Si, X. (2022). Plant-frugivore network simplification under habitat fragmentation leaves a small core of interacting generalists. Communications Biology, 5(1), 1214. 10.1038/s42003-022-04198-8

Marini, M. Â. (1992). Foraging Behavior and Diet of the Helmeted Manakin. The Condor (Los Angeles, Calif.), 94(1), 151–158. 10.2307/1368804

Martins, A. E., de Loiola, P. de P., Pareja-Bonija, D., & Morellato, L. P. C. (2025a). Climate-induced shifts in long-term tropical tree reproductive phenology: Insights from species dependent on and independent of biotic pollination. Functional Ecology. 10.1111/1365-2435.70090

Martins, A. E., de Loiola, P. de P., Pareja-Bonija, D., & Morellato, L. P. C. (2025b). Climate-induced shifts in long-term tropical tree reproductive phenology: Insights from species dependent on and independent of biotic pollination. Functional Ecology. 10.1111/1365-2435.70090

Mota, F. M. M., Heming, N. M., Morante-Filho, J. C., & Talora, D. C. (2022). Climate change is expected to restructure forest frugivorous bird communities in a biodiversity hot-point within the Atlantic Forest. Diversity and Distributions, 28(12), 2886–2897. 10.1111/ddi.13602

Naimi, B., & Araújo, M. B. (2016). Sdm: A reproducible and extensible R platform for species distribution modelling. Ecography, 39(4), 368–375. 10.1111/ecog.01881

Nascimento, E. L. M. do, Velazco, S. J. E., Ramos, F. M., Ramos, R. G., Soterroni, A. C., & Tessarolo, G. (2025). Climate change and feeble governance threaten the endangered endemic Cerrado flora in Brazil. Perspectives in Ecology and Conservation, 23(4), 290–299. 10.1016/j.pecon.2025.08.007

Nuñez, T. A., Prugh, L. R., & Hille Ris Lambers, J. (2023). Animal-mediated plant niche tracking in a changing climate. Trends in Ecology & Evolution, 38(7), 654–665. 10.1016/j.tree.2023.02.005

Owens, H. L., Campbell, L. P., Dornak, L. L., Saupe, E. E., Barve, N., Soberón, J., Ingenloff, K., Lira-Noriega, A., Hensz, C. M., Myers, C. E., & Peterson, A. T. (2013). Constraints on interpretation of ecological niche models by limited environmental ranges on calibration areas. Ecological Modelling, 263, 10–18. 10.1016/j.ecolmodel.2013.04.011

Parmesan, C. (2006). Ecological and evolutionary responses to recent climate change. Annual Review of Ecology, Evolution, and Systematics, 37, 637–669. 10.1146/annurev.ecolsys.37.091305.110100

Parr, C. L., Lehmann, C. E. R., Bond, W. J., Hoffmann, W. A., & Andersen, A. N. (2014). Tropical grassy biomes: misunderstood, neglected, and under threat. Trends in Ecology & Evolution, 29(4), 205–213. 10.1016/j.tree.2014.02.004

Pearson, R. G., & Dawson, T. P. (2003). Predicting the impacts of climate change on the distribution of species: Are bioclimate envelope models useful? Global Ecology and Biogeography, 12(5), 361–371. 10.1046/j.1466-822X.2003.00042.x

Pebesma, E. (2018). Simple Features for R: Standardized Support for Spatial Vector Data. The R Journal, 10(1), 439. 10.32614/RJ-2018-009

Pérez-Harguindeguy, N., Díaz, S., Garnier, E., Lavorel, S., Poorter, H., Jaureguiberry, P., Bret-Harte, M. S., Cornwell, W. K., Craine, J. M., Gurvich, D. E., Urcelay, C., Veneklaas, E. J., Reich, P. B., Poorter, L., Wright, I. J., Ray, P., Enrico, L., Pausas, J. G., de Vos, A. C., … Cornelissen, J. H. C. (2013). New handbook for standardised measurement of plant functional traits worldwide. Australian Journal of Botany, 61(3), 167–234. 10.1071/BT12225

Peterson, T., Soberón, J., Pearson, R. G., Anderson, R. P., Martínez-Meyer, E., Nakamura, M., & Araújo, M. B. (2011). Ecological Niches and Geographic Distributions (MPB-49). Princeton University Press. 10.23943/princeton/9780691136868.001.0001

Pinheiro, J., & Bates, D. (1999). nlme: Linear and Nonlinear Mixed Effects Models. In CRAN: Contributed Packages. 10.32614/CRAN.package.nlme

Poggio, L., de Sousa, L. M., Batjes, N. H., Heuvelink, G. B. M., Kempen, B., Ribeiro, E., & Rossiter, D. (2021). SoilGrids 2.0: producing soil information for the globe with quantified spatial uncertainty. SOIL, 7(1), 217–240. 10.5194/soil-7-217-2021

Poisot, T., Canard, E., Mouillot, D., Mouquet, N., & Gravel, D. (2012). The dissimilarity of species interaction networks. Ecology Letters, 15(12), 1353–1361. 10.1111/ele.12002

Quintero, I., & Wiens, J. J. (2013). Rates of projected climate change dramatically exceed past rates of climatic niche evolution among vertebrate species. Ecology Letters, 16(8), 1095–1103. 10.1111/ele.12144

Radchuk, V., Reed, T., Teplitsky, C., van de Pol, M., Charmantier, A., Hassall, C., Adamík, P., Adriaensen, F., Ahola, M. P., Arcese, P., Miguel Avilés, J., Balbontin, J., Berg, K. S., Borras, A., Burthe, S., Clobert, J., Dehnhard, N., de Lope, F., Dhondt, A. A., … Kramer-Schadt, S. (2019). Adaptive responses of animals to climate change are most likely insufficient. Nature Communications, 10(1). 10.1038/s41467-019-10924-4

Ratter, J. A., Bridgewater, S., & Ribeiro, J. F. (2003). Analysis of the floristic composition of the Brazilian Cerrado vegetation III: comparison of the woody vegetation of 376 areas. Edinburgh Journal of Botany, 60(1), 57–109. 10.1017/S0960428603000064

Rossi, L. C., Emer, C., Charles Lees, A., Berenguer, E., Barlow, J., Ferreira, J., França, F. M., Ramos, Y. G., Tavares, P., & Aurelio Pizo, M. (2025). Anthropogenic disturbances simplify frugivory interactions in Amazonia. Oikos, 2025(7). 10.1002/oik.10831

Rousset, F., Ferdy, J.-B., & Courtiol, A. (2013). spaMM: Mixed-Effect Models, with or without Spatial Random Effects. In CRAN: Contributed Packages. 10.32614/CRAN.package.spaMM

Royle, J. A., Chandler, R. B., Yackulic, C., & Nichols, J. D. (2012). Likelihood analysis of species occurrence probability from presence-only data for modelling species distributions. Methods in Ecology and Evolution, 3(3), 545–554. 10.1111/j.2041-210X.2011.00182.x

Sales, L., Culot, L., & Pires, M. M. (2020). Climate niche mismatch and the collapse of primate seed dispersal services in the Amazon. Biological Conservation, 247(April), 108628. 10.1016/j.biocon.2020.108628

Sales, L., Galetti, M., & Pires, M. (2020). Climate and land-use change will lead to a faunal “savannization” on tropical rainforests. Global Change Biology, 26(12), 7036–7044. 10.1111/gcb.15374

Sandor, M. E., Aslan, C. E., Pejchar, L., & Bronstein, J. L. (2021). A Mechanistic Framework for Understanding the Effects of Climate Change on the Link Between Flowering and Fruiting Phenology. Frontiers in Ecology and Evolution, 9. 10.3389/fevo.2021.752110

Schleuning, M., Neuschulz, E. L., Albrecht, J., Bender, I. M. A., Bowler, D. E., Dehling, D. M., Fritz, S. A., Hof, C., Mueller, T., Nowak, L., Sorensen, M. C., Böhning-Gaese, K., & Kissling, W. D. (2020). Trait-Based Assessments of Climate-Change Impacts on Interacting Species. Trends in Ecology & Evolution, 35(4), 319–328. 10.1016/j.tree.2019.12.010

Simon, L. M., Oliveira, G. de, Barreto, B. de S., Nabout, J. C., Rangel, T. F. L. V. B., & Diniz-Filho, J. A. F. (2013). Effects of global climate changes on geographical distribution patterns of economically important plant species in cerrado. Revista Árvore, 37(2), 267–274. 10.1590/S0100-67622013000200008

Strassburg, B. B. N., Brooks, T., Feltran-Barbieri, R., Iribarrem, A., Crouzeilles, R., Loyola, R., Latawiec, A. E., Oliveira Filho, F. J. B., De Scaramuzza, C. A. M., Scarano, F. R., Soares-Filho, B., & Balmford, A. (2017). Moment of truth for the Cerrado hotspot. In Nature Ecology and Evolution (Vol. 1, Number 4). Nature Publishing Group. 10.1038/s41559-017-0099

Thébault, E., & Fontaine, C. (2010). Stability of Ecological Communities and the Architecture of Mutualistic and Trophic Networks. Science, 329(5993), 853–856. 10.1126/science.1188321

Thuiller, W. (2024). Ecological niche modelling. Current Biology, 34(6), R225–R229. 10.1016/j.cub.2024.02.018

Timóteo, S., Ramos, J. A., Vaughan, I. P., & Memmott, J. (2016). High Resilience of Seed Dispersal Webs Highlighted by the Experimental Removal of the Dominant Disperser. Current Biology, 26(7), 910–915. 10.1016/j.cub.2016.01.046

Tobias, J. A., Sheard, C., Pigot, A. L., Devenish, A. J. M., Yang, J., Sayol, F., Neate-Clegg, M. H. C., Alioravainen, N., Weeks, T. L., Barber, R. A., Walkden, P. A., MacGregor, H. E. A., Jones, S. E. I., Vincent, C., Phillips, A. G., Marples, N. M., Montaño-Centellas, F. A., Leandro-Silva, V., Claramunt, S., … Schleuning, M. (2022). AVONET: morphological, ecological and geographical data for all birds. Ecology Letters, 25(3), 581–597. 10.1111/ele.13898

Vázquez, D. P., & Aizen, M. A. (2004). Asymmetric specialization: A pervasive feature of plant-pollinator interactions. Ecology, 85(5), 1251–1257. 10.1890/03-3112

Vázquez, D. P., Blüthgen, N., Cagnolo, L., & Chacoff, N. P. (2009). Uniting pattern and process in plant–animal mutualistic networks: a review. Annals of Botany, 103(9), 1445–1457. 10.1093/aob/mcp057

Velazco, S. J. E., Villalobos, F., Galvão, F., & De Marco Júnior, P. (2019). A dark scenario for Cerrado plant species: Effects of future climate, land use and protected areas ineffectiveness. Diversity and Distributions, 25(4), 660–673. 10.1111/ddi.12886

Vincenty, T. (1975). Direct and inverse solutions of geodesics on the ellipsoid with applications of nested equations. Survey Review, 23(176), 88–93. 10.1179/sre.1975.23.176.88

Vizentin-Bugoni, J., & Maruyama, P. K. (2023). To rewire or not to rewire: To what extent rewiring to surviving partners can avoid extinction? Journal of Animal Ecology, 92(9), 1676–1679. 10.1111/1365-2656.13972

Vizentin-Bugoni, J., Maruyama, P. K., & Sazima, M. (2014). Processes entangling interactions in communities: forbidden links are more important than abundance in a hummingbird–plant network. Proceedings of the Royal Society B: Biological Sciences, 281(1780), 20132397. 10.1098/rspb.2013.2397

Vizentin-Bugoni, J., Sperry, J. H., Kelley, J. P., Foster, J. T., Drake, D. R., Case, S. B., Gleditsch, J. M., Hruska, A. M., Wilcox, R. C., & Tarwater, C. E. (2022). Mechanisms underlying interaction frequencies and robustness in a novel seed dispersal network: Lessons for restoration. Proceedings of the Royal Society B: Biological Sciences, 289(1982). 10.1098/rspb.2022.1490

Vizentin-Bugoni, J., Tarwater, C. E., Foster, J. T., Drake, D. R., Gleditsch, J. M., Hruska, A. M., Kelley, J. P., & Sperry, J. H. (2019). Structure, spatial dynamics, and stability of novel seed dispersal mutualistic networks in Hawai1i. Science, 364(6435), 78–82. 10.1126/science.aau8751

Wilman, H., Belmaker, J., Simpson, J., de la Rosa, C., Rivadeneira, M. M., & Jetz, W. (2014). EltonTraits 1.0: Species-level foraging attributes of the world’s birds and mammals. Ecology, 95(7), 2027–2027. 10.1890/13-1917.1

Zaca, W., Silva, W. R., & Pedroni, F. (2006). Diet of the Rusty-margined Guan (penelope Superciliaris) in an Diet of the Rusty-margined Guan (penelope Superciliaris) in an Altitudinal Forest Fragment of Southeastern Brazil Altitudinal Forest Fragment of Southeastern Brazil. Ornitología Neotropical Ornitología Neotropical, 17(3), 5.

