## Supplementary informations for "Climate change is predicted to simplify seed dispersal networks in the Cerrado"

- **Sampling of plant–frugivore interactions from the compiled dataset**

Plant–frugivore interaction data used in this study were obtained from the dataset published by (Campagnoli et al., 2025). In that study, interactions were recorded monthly along six 100 × 10 m transects using three complementary sampling methods: focal observations, faecal samples from birds captured with mist nets, and videos from camera traps. Sampling effort ranged from 41.5 to 95.35 h per transect for focal observations, 302.75 to 453.69 h across transects for mist nets, and 141.07 to 2938.37 h per plant species for camera traps.

In the original study, focal observations consisted of walking along each transect while searching for fruiting plants and recording plant–frugivore interactions. Whenever an interaction between animals and fruiting plants was observed, the observer paused the survey to record the interaction before continuing the transect walk. In addition, mist nets were installed in four transects and sampled once per month when logistics allowed. During each sampling event, five mist nets were used: three measuring 3 m in height and 7 m in length (16 mm mesh), and two measuring 2.5 m in height and 7 m in length (20 mm mesh). Nets were opened for 2–3 days and checked every 30 min from dawn to dusk, with a pause from 11:30 to 14:30 to avoid peak summer heat. Captured birds were placed in cotton bags for approximately 20–30 min or until defecation occurred. Bags were then inspected for seeds, which were subsequently identified to determine plant–frugivore interactions. Camera traps were installed monthly in transects where fruiting plants were available. Because only four cameras were available, they were rotated among transects. Cameras were positioned in front of fruiting plants whenever possible, typically selecting individuals with the highest number of ripe fruits to maximize the probability of recording frugivory events. Cameras recorded 30 s videos at 2 min intervals. An interaction event was recorded when a bird removed seeds from the plant (focal observations and camera traps) and/or when seeds of a given plant species were found in a bird faecal sample (captures with mist nets).

Empirical plant–frugivore interaction frequencies were calculated combining the three sampling methods data, following the procedure described by Campagnoli et al. (2025). Briefly, interaction frequencies from each sampling method were organized into separate interaction networks, weighted by the pooled number of interactions recorded across transects. Sampling completeness was then assessed for each network, and interaction frequencies were standardized using the grand total standardization method, following Quintero et al., 2022. This approach converts interaction frequencies recorded by each sampling method into the probability that a given pairwise interaction occurs among all recorded interactions. First, interaction frequencies were weighted by sampling effort by dividing the number of recorded interactions by the number of sampling hours associated with each method. For focal observations and mist-net captures, interaction frequencies were divided by the total sampling hours across all transects, whereas for camera traps they were divided by the total sampling hours per plant species. Second, values in each interaction matrix were weighted by the total number of interactions recorded per hour for each sampling method. Third, matrices were weighted according to the sampling completeness of the respective method. Finally, the weighted matrices were combined by calculating the average value for each pairwise interaction, resulting in a single empirical matrix of plant-frugivore interaction frequencies. This matrix was used to compare empirical interaction frequencies with the predicted interaction probability matrix based on geographic range overlap, temporal overlap, and trait matching of interacting species.

- Species occurrence sources

Species occurrences were collected from global biodiversity aggregation infrastructures, including the Global Biodiversity Information Facility (GBIF; www.gbif.org), iNaturalist (www.inaturalist.org), the Berkeley Ecoinformatics Engine (Ecoengine; www.ecoengine.berkeley.edu), and the Botanical Information and Ecological network (BIEN; https://bien.nceas.ucsb.edu). Taxonomy and nomenclature for frugivores followed the most recent IUCN Red List consensus (search date: May

2025), whereas for plants we followed the World Flora Online (WFO; https://wfoplantlist.org/) (search date: May 2025).

- **Data cleaning**

We carefully cleaned the occurrence data using the R package 'CoordinateCleaner' (Zizka et al., 2019). This involved removing duplicates and incomplete coordinate records, records located on centroids of municipal and political polygons, oceans, museums, herbaria, and those located more than 200 km outside the border of the IUCN species-specific geographic range (only for frugivores). Furthermore, occurrences with georeferencing uncertainty >=1 km and recorded before 1970 were excluded to match the spatial-temporal resolution of the environmental data. To minimize the impact of spatial autocorrelation, we spatially thinned occurrences less than 1 km from each other using the R package 'spThin' (Aiello-Lammens et al., 2015).

- **Ecological Niche Models**

We fitted ecological niche models (ENMs) to establish a statistical relationship between the species occurrences and the environmental variables (climate and soil variables). To assess model performance, we used 75% of the occurrence records for calibration and 25% for model evaluation. This subsampling procedure was repeated 15 times for each algorithm and species, resulting in 45 models for each species. As accuracy metrics, we used the true skill statistic (TSS) and the area under the receiver operating characteristic curve (AUC). The TSS metric was also used to determine a threshold to convert continuous habitat suitability predictions into binary maps (Table S1). This threshold was identified as the one that maximized the TSS for each species, ensuring optimal differentiation between suitable and unsuitable habitats by maximizing the sum of sensitivity and specificity (Liu et al., 2005). Subsequently, we implemented a 'weighted' method in the R package ‘sdm’, version 1.2 (Naimi & Araújo, 2016) to create ensembles of all models for each species, where the predictions of individual models are combined into a single consensus prediction. This weighted approach ensures that models with high predictive performance, as determined by the TSS metric, contribute more to the ensemble than models with low performance (Araújo & New, 2007). For each species, we defined a specific study area/background to fit the ENMs. These areas were delimited by a bounding box using the extreme longitude and latitude coordinates from the occurrence records (Sales et al., 2021). We further extended the bounding box by an additional 10 degrees in each direction to encompass regions potentially accessible to the species through dispersal over 100 years (Barve et al., 2011; Sales et al., 2021). We adjusted all environmental layers to match each species-specific background area and sampled the same number of pseudo-absences as species presence records.

For each species, we checked multicollinearity of the environmental variables using a hybrid approach implemented in the ‘vifcor' function from the sdm R package (Naimi & Araújo, 2016). The function calculates the variance inflation factor (VIF) for a set of environmental variables and identifies pairs with a correlation coefficient greater than a certain threshold (here $r$=0.60). From these pairs, the variable with the higher VIF is excluded. This process resulted in six to eight predictor variables for each species. Supplementary Table S1 lists all the environmental predictors used in each species' ENM.

### References


Aiello-Lammens, M. E., Boria, R. A., Radosavljevic, A., Vilela, B., & Anderson, R. P. (2015). spThin: An R package for spatial thinning of species occurrence records for use in ecological niche models. *Ecography*, *38*(5), 541–545. https://doi.org/10.1111/ecog.01132

Araújo, M. B., & New, M. (2007). Ensemble forecasting of species distributions. *Trends in Ecology & Evolution* , *22*(1), 42–47. https://doi.org/10.1016/j.tree.2006.09.010

Barve, N., Barve, V., Jiménez-Valverde, A., Lira-Noriega, A., Maher, S. P., Peterson, A. T., Soberón, J., & Villalobos, F. (2011). The crucial role of the accessible area in ecological niche modeling and species distribution modeling. *Ecological Modelling*, *222*(11), 1810–1819. https://doi.org/10.1016/j.ecolmodel.2011.02.011

Campagnoli, M., Christianini, A., & Peralta, G. (2025). Plant and frugivore species characteristics drive frugivore contributions to seed dispersal effectiveness in a hyperdiverse community. *Functional Ecology*, *39*(1), 238–253. https://doi.org/10.1111/1365-2435.14697

Liu, C., Berry, P. M., Dawson, T. P., & Pearson, R. G. (2005). Selecting thresholds of occurrence in the prediction of species distributions. *Ecography*, *28*(3), 385–393. https://doi.org/10.1111/j.0906-7590.2005.03957.x

Naimi, B., & Araújo, M. B. (2016). Sdm: A reproducible and extensible R platform for species distribution modelling. *Ecography*, *39*(4), 368–375. https://doi.org/10.1111/ecog.01881

Quintero, E., Isla, J., & Jordano, P. (2022). Methodological overview and data-merging approaches in the study of plant–frugivore interactions. *Oikos*, *2022*(2). https://doi.org/10.1111/oik.08379

Sales, L., Kissling, W. D., Galetti, M., Naimi, B., & M. Pires, M. (2021). Climate change reshapes the eco-evolutionary dynamics of a Neotropical seed dispersal system. *Global Ecology and Biogeography*, *30*(5), 1129–1138. https://doi.org/10.1111/geb.13271

Zizka, A., Silvestro, D., Andermann, T., Azevedo, J., Duarte Ritter, C., Edler, D., Farooq, H., Herdean, A., Ariza, M., Scharn, R., Svantesson, S., Wengström, N., Zizka, V., &

Antonelli, A. (2019). CoordinateCleaner: Standardized cleaning of occurrence records from biological collection databases. *Methods in Ecology and Evolution*, *10*(5), 744–751. https://doi.org/10.1111/2041-210X.13152
